# Inflammation is a stronger predictor of psychological trauma exposure than behavior in repeated social defeat

**DOI:** 10.1101/2020.05.18.102129

**Authors:** Safwan K. Elkhatib, Cassandra M. Moshfegh, Gabrielle F. Watson, Adam J. Case

## Abstract

Post-traumatic stress disorder (PTSD) is a psychiatric illness that results in an increased risk for a variety of inflammatory diseases. The exact etiology of this increased risk in unknown, and thus, several animal models have been developed to investigate the neuroimmune interactions of PTSD. Repeated social defeat stress (RSDS) is an established preclinical model of psychological trauma that recapitulates certain behavioral and inflammatory aspects of human PTSD. Furthermore, RSDS has been utilized to subgroup animals into susceptible and resilient populations based on one specific behavioral phenotype (*i.e*., social interaction). Herein, we conducted an extensive investigation of circulating inflammatory proteins after RSDS, and found significant elevations in various cytokines and chemokines after exposure to psychological trauma. When categorizing animals into either susceptible or resilient populations based on social interaction, we found no inflammatory or other behavioral differences between these subgroups. Furthermore, assessment of associations between all detectable inflammatory proteins and behavioral outputs found no significant correlation between social interaction parameters and inflammation. In contrast, we identified a panel of 5 circulating inflammatory proteins that showed significant associations with elevated zero maze parameters. Strikingly, these 5 circulating inflammatory proteins displayed a stronger predictive ability of psychological trauma exposure compared to any behavioral outcome. These findings provide new insights into inflammatory markers associated with RSDS, and their ability to predict psychological trauma exposure more robustly than commonly utilized behavioral paradigms.

**Highlights:**

- Repeated social defeat stress (RSDS) reproducibly produces peripheral inflammation
- Peripheral inflammation is not coupled to social interaction testing parameters
- Susceptible and resilient categorization does not reflect peripheral inflammation
- Anxiety-like behavioral parameters are linked to peripheral inflammation in RSDS
- Peripheral inflammation is more predictive of trauma than behavior in RSDS

## Introduction

Post-traumatic stress disorder (PTSD) is a devastating psychiatric illness characterized by exposure to a traumatic event, followed by flashbacks, avoidance, affective changes, and hyperarousal.^1^ Interestingly, only about 20% of those that experience a traumatic event develop clinical PTSD, which demonstrates trauma exposure is necessary, but not sufficient alone, to cause PTSD.^2^ Given this, there has been a continued interest in determining which psychological or biological factors could determine susceptibility or resilience to PTSD.^3, 4^ Understanding that PTSD patients notably face an increased risk for a number of inflammation-driven diseases,^5–9^ investigations into the inflammatory contributions to PTSD have recently garnered significant attention (reviewed exceptionally by Sumner *et al*.).^10^

In attempts to elucidate the pathophysiology of PTSD, many preclinical rodent models have emerged.^11, 12^ Repeated social defeat stress (RSDS) is a well-characterized murine model that relies upon the aggressive and territorial nature of retired breeder CD-1 mice,^13^ while successfully recapitulating several behavioral and inflammatory aspects of PTSD.^11, 12^ Furthermore, this model has been reported to produce both susceptible and resilient populations as defined by a single parameter on the social interaction behavioral test (*i.e*., social interaction ratio <1.0 or ≥1.0, respectively).^13, 14^

Recent work from our group utilized the model of RSDS to examine the autonomic and redox profiles of T-lymphocytes after psychological trauma exposure.^15^ Interestingly, while we demonstrated significant and novel changes to T-lymphocyte-driven inflammation, these did not correlate well with behavior assessed by the social interaction test, despite individually showing differences between RSDS and control cohorts.^15^ With this observation, we hypothesized that peripheral inflammation and behavior are not tightly coupled following psychological trauma exposure.

In the present study, we sought to identify links between two standard RSDS behavioral tests (*i.e*., social interaction test and elevated zero maze) and peripheral inflammation using a large-scale assessment of circulating plasma inflammatory proteins following RSDS. In doing so across a large cohort of animals, we found little data to support any inflammatory differences between susceptible and resilient groups of animals determined by the social interaction ratio. Notably, we further identified a select number of circulating inflammatory proteins as better predictors of exposure to psychological trauma than either behavioral test.

## Methods

### Mice

RSDS was performed as previously described.^15^ In brief, control and RSDS animals were 8-12 week-old wild-type male mice of a C57BL/6J background (Jackson Laboratory #000664, Bar Harbor, ME, USA). Seventy-four total animals were used in this study. Standardized RSDS paradigms canonically utilize male mice only,^13^ thus biological sex as a variable was not examined herein. All experimental mice were bred in-house and group-housed (≤5 mice/cage) prior to stress induction to attenuate possible confounding physical, psychological, and social stressors. Aggressive mice used for RSDS induction were 4-8 month-old retired breeder male mice of a CD-1 background (Charles River #022, Wilmington, MA, USA). All mice were caged with corncob bedding, paper nesting material, and given access to standard chow (Teklad Laboratory Diet #7012, Harlan Laboratories, Madison, WI, USA) and water ad libitum. Experimental mice were sacrificed by intraperitoneal injection of pentobarbital (150 mg/kg, Fatal Plus, Vortech Pharmaceuticals, Dearborn, MI, USA). Daily RSDS and terminal euthanasia occurred between 07:00 and 09:00 to eliminate the known effects of circadian rhythm on inflammation. Experimental mice were randomized to control or RSDS cohorts, and all efforts were made to blind experimenters during biological assays and data analysis. All procedures were reviewed and approved by the University of Nebraska Medical Center Institutional Animal Care and Use Committee.

### RSDS

Stress induction was accomplished through a modified RSDS paradigm.^15^ Briefly, retired breeder CD-1 mice were first pre-screened and selected for aggressive behavior.^13^ After singly inhabiting cages for 3 days, experimental mice were introduced into these cages for physical confrontation for 5 minutes. For the remaining 24 hours, experimental and aggressor mice were co-housed but separated by a transparent barrier. The processes was repeated daily with a new CD-1 aggressive mouse for 10 consecutive days. Control mice were similarly pair-housed sans daily social defeat sessions. All experimental mice were behavior tested on day 11. On day 12, all mice were sacrificed for biological analysis. While animals with visual wounding (>1cm)^13^ or lameness due to stress induction are excluded from further study, none of the 74 animals utilized herein met this threshold for exclusion.

### Behavioral Testing

All behavior testing was conducted as previously described.^15^ For social interaction testing, an open field (40 x 40 cm, Noldus Information Technology, Leesburg, VA, USA) was outfitted with a transparent enclosure with mesh caging (6.5 x 10 cm). All mice were first tested in the open field with an empty enclosure, and then subsequently tested with a novel CD-1 mouse present within the enclosure. A ratio of the time spent in proximity to the enclosure (interaction zone) or distal corners of the maze in the presence versus absence of a novel CD-1 mouse defines the social interaction ratio and corner zone ratio, respectively.^13, 14^ Each run lasted 2.5 minutes, and all sessions were recorded and digitally analyzed by Noldus Ethovision XT 13 software (Leesburg, VA, USA). Susceptible and resilient populations were defined by social interaction ratio <1 or ≥1, respectively.

For elevated zero maze testing, all experimental mice were introduced into a circular elevated maze (50 cm diameter, 5 cm track width, Noldus Information Technology, Leesburg, VA, USA) with 50% enclosed (20 cm wall height) and 50% open arenas. Mice were allowed to freely explore the novel environment for 5 minutes. All sessions recorded and digitally analyzed by Noldus Ethovision XT 13 software (Leesburg, VA, USA).

### Plasma Inflammatory Protein Measurement

Blood was obtained by cardiac puncture immediately after sacrifice using ethylenediaminetetraacetic acid (EDTA) as an anticoagulant. Plasma was separated by centrifugation, and stored at −80°C until use. Plasma inflammatory protein measurements were obtained by utilizing the Meso Scale Discovery V-Plex Mouse Cytokine 29-plex kit (#K15267D, Rockville, MD, USA). Meso Scale Discovery V-plex plates are tightly validated by the manufacturer to avoid antigenic cross-reactivity. The following inflammatory proteins were measured: IFN-γ, IL-1β, IL-2, IL-4, IL-5, IL-6, IL-9, IL-10, IL-12p70, IL-15, IL-16, IL-17A, IL-17C, IL-17F, IL-17A/F, IL-17E/IL-25, IL-21, IL-22, IL-23, IL-27p28/IL-30, IL-31, IL-33, CXCL10, CXCL1, CCL2, CCL3, CXCL2, CCL20, TNF-α. All experiments were conducted per the manufacturer’s instructions, with samples run on Meso Scale Discovery Quickplex SQ 120 and analyzed using the Mesoscale Discovery Workbench software.

### Statistics

A total of 74 animals (35 control, 39 RSDS) were utilized for the following studies. Associations between behavioral assessments and inflammatory proteins were assessed by nonparametric spearman correlation with two-tailed confidence intervals. Familywise error rate for the correlation matrix was conservatively corrected with the Bonferroni procedure for 20 inferential hypotheses (16 detectable inflammatory proteins and 4 behavioral outputs), with statistical significance thus established at p<0.0025 (α/20 hypotheses tested). Receiver-operating characteristics (ROC) curves were assessed for classification ability by the area under the curve (AUC), with predicted probability for each subject set at 0.5. Goodness-of-fit was assessed by likelihood ratio test.

For inflammatory protein measurements, individual data are presented, with summary data displayed as mean ± SEM. Each data set was assessed for normality utilizing the Shapiro-Wilk test, followed by appropriate inferential hypothesis testing. For all inflammatory protein and behavioral measurements, Mann-Whitney U test was used to assess differences between control and RSDS. Susceptible and resilient are subgroups of RSDS, and thus statistical tests were only conducted between the two subgroups. All statistical analyses were completed in GraphPad Prism 8.4.2 (San Diego, CA, USA).

## Results

### Susceptibility and resilience by social interaction test does not differentiate inflammatory outcomes

After RSDS, all mice were examined for behavioral changes by social interaction and elevated zero maze tests. Social interaction and corner zone ratios showed significant and non-significant differences between control and stress, respectively (p=0.0370, p=0.0754; **Figure 1**). Elevated zero maze parameters—distance moved and time in the open arm—showed significant and robust differences after RSDS (p<0.0001, p<0.0001; **Figure 1**). When RSDS animals were subdivided into susceptible and resilient groups by social interaction ratio (susceptible = SI ratio <1; resilient = SI ratio ≥1), there were no differences between these subgroups across the corner zone ratio, distance moved, or time in open (p=0.0870, p=0.1895, p=0.0907, respectively; **Figure 1**).

**Figure 1.**
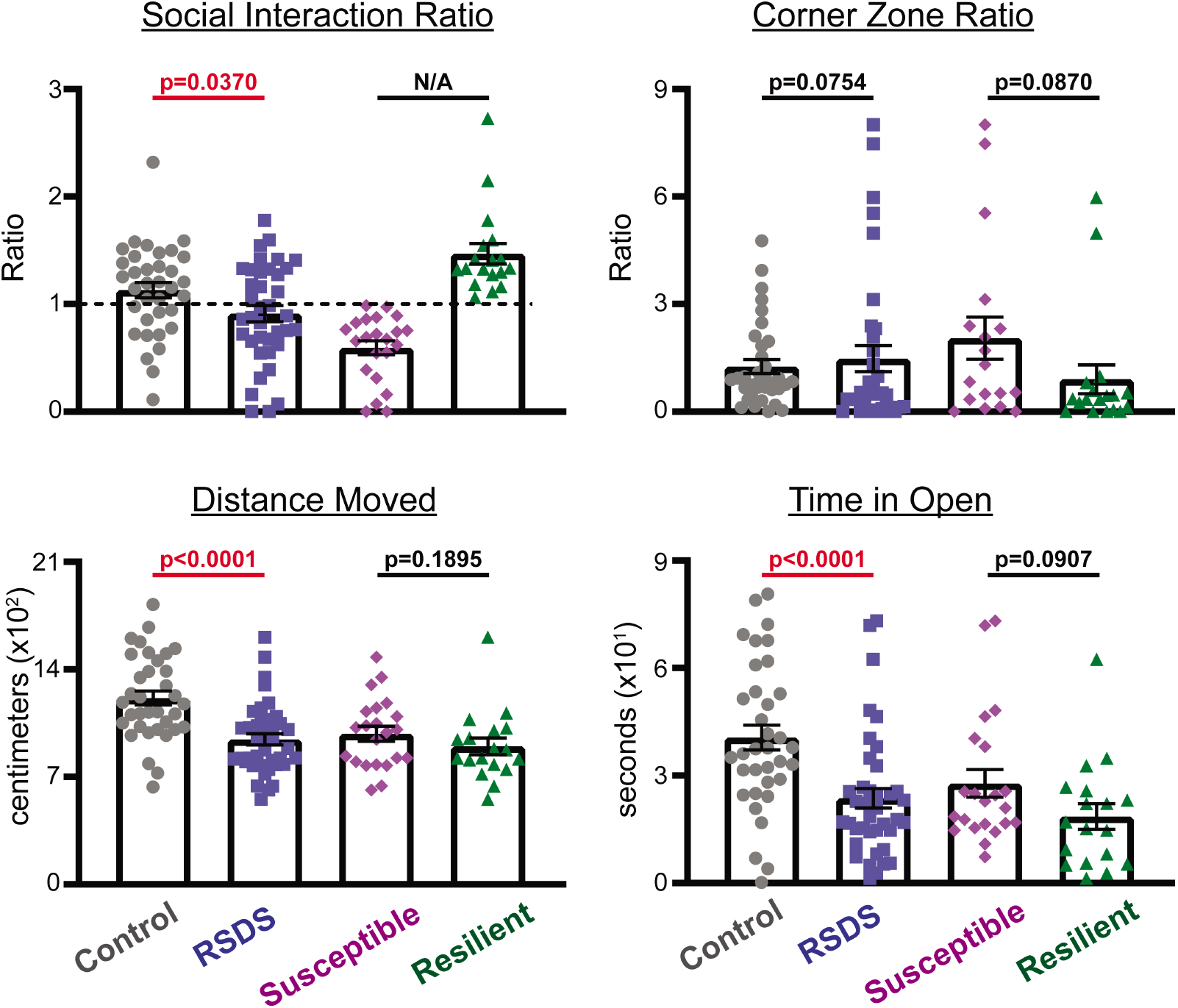
Susceptible and resilient subgrouping as determined by social interaction ratio does not reflect additional behavioral phenotypes. Mice were run through the 10-day RSDS paradigm followed by social interaction test (upper panels) and elevated zero maze (lower panels) behavioral testing. RSDS mice were categorized into susceptible or resilient subgroups by a social interaction ratio <1.0 or ≥1.0, respectively (*upper left*). N = 35 control, 39 RSDS. Statistical significance by parametric Student’s t-test or nonparametric Mann-Whitney U test where appropriate.

With peripheral inflammation as our primary biological outcome, we then investigated shifts in circulating inflammatory protein levels following RSDS. Of the 29 inflammatory proteins analyzed, 16 were found to be above the lower limit of detection: IFN-γ, IL-10, IL-16, IL-17A, IL-1β, IL-2, IL-22, IL-5, IL-6, CXCL10, CXCL1, CCL2, CCL3, CXCL2, CCL20, and TNF-α. Furthermore, 8 of these proteins showed significant differences between control and RSDS animals after correction for multiple analyses (**Figure 2**). Next, we further grouped the inflammatory proteins from RSDS animals into susceptible and resilient groups as determined by the social interaction ratio. However, no significant differences in peripheral inflammation were detected between susceptible and resilient groups (**Figure 2**). Overall, RSDS results in complex and robust changes to peripheral inflammation. However, these changes do not group into susceptible or resilient populations when separated by social interaction ratio.

**Figure 2.**
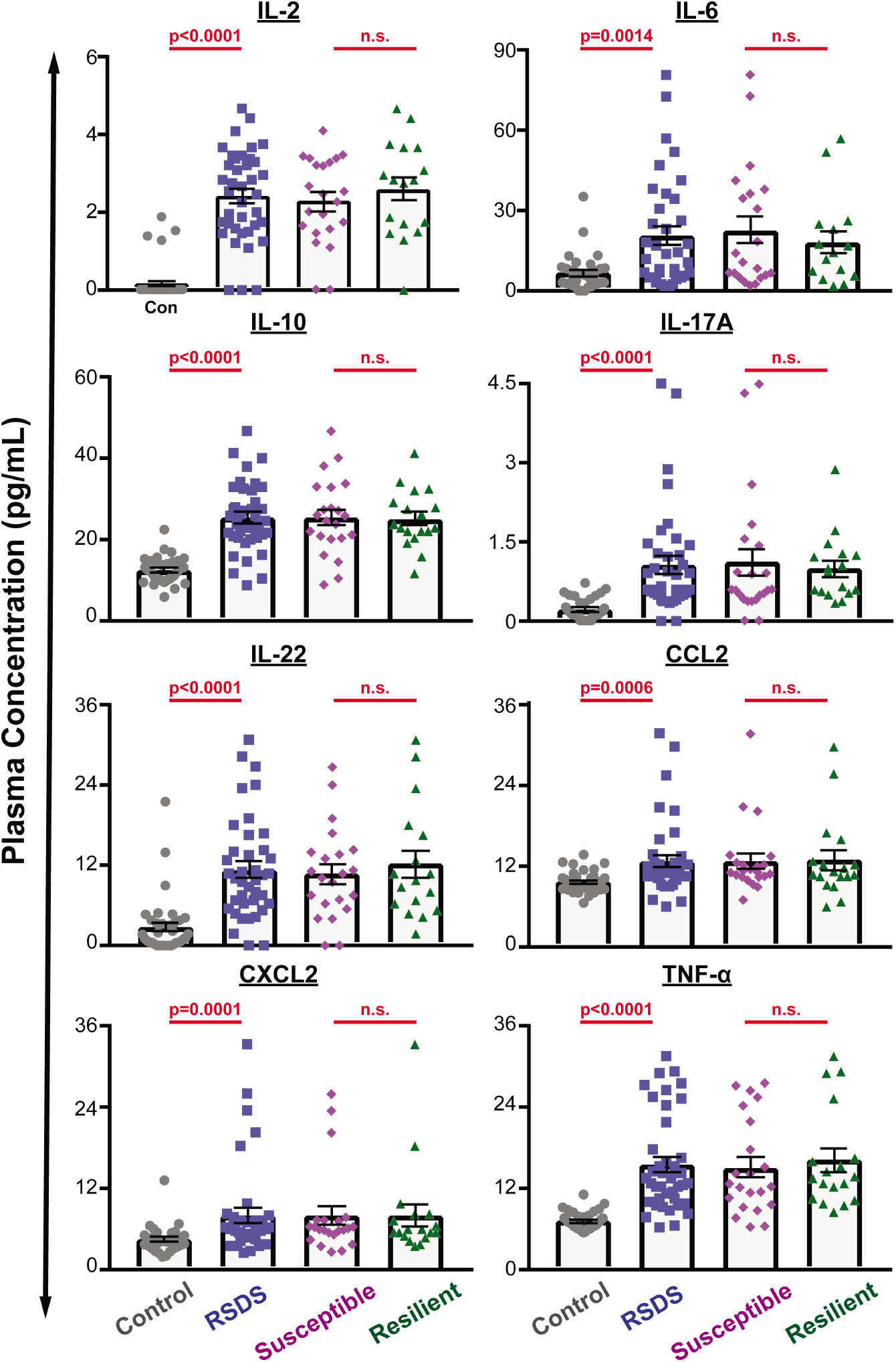
Susceptible and resilient subgrouping as determined by social interaction ratio does not reflect peripheral inflammation. Mice were run through the 10-day RSDS paradigm followed by behavioral testing and plasma extraction. RSDS mice were categorized into susceptible or resilient subgroups by a social interaction ratio <1.0 or ≥1.0, respectively. Circulating levels of 29 cytokines were assessed by Meso Scale Discovery multiplex analysis; 16 were above the limit of detection, and 8 showed significant differences between control and RSDS animals. N = 35 control, 39 RSDS. Statistical significance by nonparametric Mann-Whitney U test.

### Selected circulating cytokines predict trauma exposure more accurately than behavioral assessments

In order to conduct a more comprehensive examination into the associations between common behavioral assessments and circulating inflammation, we generated a correlation matrix between the 16 detectable inflammatory proteins and the aforementioned behavioral parameters in all animals. We found that social interaction test parameters (*i.e*., social interaction ratio and corner zone ratio) did not significantly correlate with any detectable inflammatory protein (**Figure 3A**). However, elevated zero maze outputs (*i.e*., distance moved and time in open) associated significantly with 5 inflammatory proteins (**Figure 3A**): IL-10 (R= −0.443, −0.448), IL-17A (R= −0.420, −0.466), IL-2 (= - 0.501, −0.484), IL-22 (R= −0.451, −0.445), and TNF-α (R= −0.510, −0.563).

Next, multiple logistic regression was utilized to assess the ability of these behavioral and inflammatory parameters to predict psychological trauma exposure. From this regression model, ROC curves were generated. While the two social interaction test parameters did not result in significant predictive ability (AUC=0.5632, p=0.3509, G^2^ log likelihood ratio=4.86, p=0.088; **Figure 3B**), the two elevated zero maze parameters were able to significantly predict psychological trauma exposure (AUC=0.786, p<0.0001, G^2^ log likelihood ratio=19.31, p<0.0001; **Figure 3B**). Unexpectedly, the combination of the 5 significant inflammatory proteins (IL-10, IL-17A, IL-2, IL-22, TNF-α) that correlated with elevated zero maze parameters were able to predict psychological trauma exposure more than either behavioral test alone or in combination (AUC=0.9706, p<0.0001, G^2^ log likelihood ratio=77.41, p<0.0001; **Figure 3B**). Overall, we demonstrate that elevated zero maze parameters associate more strongly with peripheral inflammation than the social interaction test after RSDS. Additionally, psychological trauma exposure is better predicted by select circulating inflammatory measures than either social interaction test or elevated zero maze.

**Figure 3.**
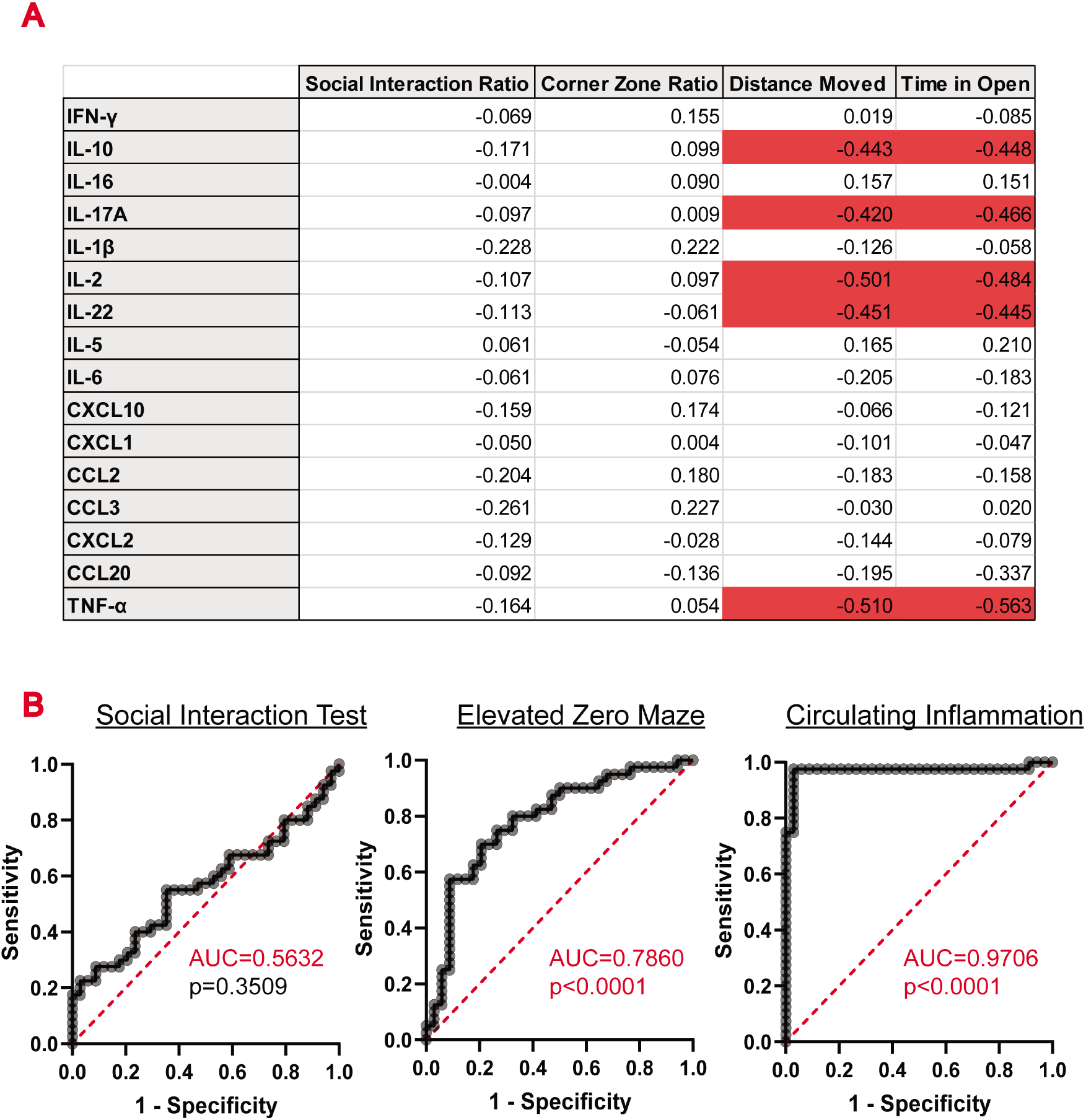
A specific subset of peripheral inflammatory proteins is more predictive of RSDS exposure than behavioral phenotypes. Mice were run through the 10-day RSDS paradigm followed by behavioral testing, plasma extraction, and inflammation assessment by Meso Scale Discovery multiplex analysis. **A**. Correlation matrix of behavioral parameters and circulating inflammatory proteins. Values listed are nonparametric Spearman R coefficients (two-tailed test), with significant associations highlighted (red). **B**. ROC curves were generated for social interaction test (social interaction and corner zone ratios together), elevated zero maze (distance moved and time in open together), and significant inflammatory parameters (IL-2, IL-10, IL-17A, IL-22, and TNFα together). Prediction ability was calculated by area under curve, with p-value calculated by a two-tailed test utilizing a predicted probability of each subject at 0.5.

## Discussion

In the current study, we confirm that RSDS results in specific behavioral and peripheral inflammatory alterations. However, when categorizing RSDS into susceptible and resilient groups indexed by the social interaction ratio, we did not observe similar grouping by other behavioral outputs. Indeed, in the original categorization of susceptible and resilient by social interaction ratio by Krishnan *et al*., the authors also conceded this grouping did not reflect anxiety-like behavior, locomotor activity, stress-induced polydipsia, or despair behavior after RSDS.^14^ While this categorization of mice has indeed uncovered biological differences between these social interaction ratio-derived populations, these findings suggest this grouping is not all-encompassing, and may be limited to only a specific behavioral manifestation that may or may not be related to human disease.

Additionally, we demonstrate herein that social interaction ratio grouping does not meaningfully divide RSDS animals by peripheral inflammatory changes. When examining 29 inflammatory proteins in circulation, we found 8 that were differentially extant in the plasma of RSDS mice. However, none of these inflammatory proteins were significantly different between susceptible and resilient groups when categorized by the social interaction ratio. This data prompts further query into the usage of social interaction ratio to determine subpopulations of susceptible and resilient from RSDS, especially when examining peripheral inflammation.

Intriguingly, by examining associations between social interaction, elevated zero maze, and peripheral inflammation, we found that 5 circulating inflammatory proteins (of the original 8 that were significantly different) were better able to predict psychological trauma exposure than either behavioral test. While highly counterintuitive, these data suggest behavioral outcomes (at least these specific tests) are not tightly linked to the exposure to psychological trauma, and more importantly, do not predict physiological changes in trauma-exposed individuals. This finding may be of significant relevance to humans exposed to psychological trauma. Aforementioned, only approximately 20% of individuals exposed to significant trauma develop PTSD. This means at approximately 80% are deemed “resilient” to the disease, however, PTSD diagnosis is only made based on psychological manifestations after trauma exposure, not physiological. Given our results, this may mean many individuals exposed to trauma display inflammatory changes independent of behavioral changes, and therefore may go undiagnosed. This finding has significant clinical implications, and suggests inflammatory investigations into trauma-exposed patients, not only those diagnosed with PTSD, is highly warranted.

Based on our findings, an important aspect for discussion is how to appropriately segment traumatized animals into susceptible and resilient populations. While the social interaction ratio provides a convenient methodology for separating RSDS mice, we have presented data herein to shown that it does not effectively mirror changes in anxiety-like behavior or peripheral inflammation. Additionally, recent work has shown that the social interaction test itself appears to involve conditioned learning based on prior exposures to an aggressive white CD-1 mice, versus true tests of antisocial behavior.^16^ These data should serve as unembellished reminders of the inherent complexity to which psychological and physiological factors might guide the development of susceptibility or resilience—even within our more reductionist preclinical models. Moreover, these data support the sentiment of recommendations by the National Institute of Mental Health *Research Domain Criteria* (RDoC),^17^ which seeks to represent all behavioral characterizations as continuous variables across multiple domains that allow for more definitive (and complex) analyses.^18^ The development of multifaceted approaches to assess the degree of psychological trauma will be paramount in further elucidation of the pathophysiology of this disease, as well as assessment of the efficacy of potential therapeutics.

While RSDS has shown robust and consistent peripheral inflammatory changes, one potentially problematic aspect of the model is the prospect of wounding. While we take care to exclude all animals with significant wounds or lameness, as previously discussed, this remains a possible confounder in RSDS and other models that incorporate physical stress. We estimate at least 75% of all animals in this study showed no visible signs of wounding, yet virtually all animals displayed a peripheral inflammatory response. These data are suggestive that wounding does not fully explain the inflammatory changes of these mice, but at this time cannot be completely ruled out. However, it should be noted that psychological trauma in humans very often is accompanied by physical trauma. This is especially vital in the forms of trauma that are most likely to result in PTSD—rape, physical assault, and serious injury— and that may also directly impact the immune system.^19^

The present study and its implications are of course not without limitations. The most crucial in its external validity is the lack of examination of sex as a biological variable. Due to the introduction of aggressive retired-breeder male mice in traditional RSDS, it cannot be used to induce psychological trauma similarly in female mice. There have been recent reports of modified female social defeat stress,^20, 21^ which—while potentially effective insofar as stress induction—vary significantly enough in approach, intensity, and duration to serve as serious confounding variables reducing internal validity. For this reason, this was not addressed in the present study. Future examination should include these behavioral and inflammatory paradigms in the context of newly developed models of female social defeat stress.

In summary, this current study provides novel findings that prompt a more nuanced approach to determination of susceptibility and resilience by use of the social interaction ratio. Further, we present a select peripheral inflammatory panel that better predicts psychological trauma exposure compared to standardized behavior tests. Together, these findings call for a more in-depth and mechanistic analysis of psychological trauma in the RSDS model, as well as translational examination of peripheral inflammation in trauma-exposed human individuals.

## Ethics

This study was carried out in accordance with the recommendations of the University of Nebraska Medical Center Institutional Animal Care and Use Committee. The protocol was approved by the University of Nebraska Medical Center Institutional Animal Care and Use Committee.

## Author Contributions

SE, CM, GW, and AC designed all research questions and studies. SE, CM, and GW conducted experiments and analyzed data. SE and AC wrote the manuscript. SE, CM, GW, and AC reviewed, edited, and approved the manuscript. AC provided experimental oversight and funding.

